# The effect of stimulus choice on an EEG-based objective measure of speech intelligibility

**DOI:** 10.1101/421727

**Authors:** Eline Verschueren, Jonas Vanthornhout, Tom Francart

## Abstract

**Objectives:** Recently an objective measure of speech intelligibility, based on brain responses derived from the electroencephalogram (EEG), has been developed using isolated Matrix sentences as a stimulus. We investigated whether this objective measure of speech intelligibility can also be used with natural speech as a stimulus, as this would be beneficial for clinical applications.

**Design:** We recorded the EEG in 19 normal-hearing participants while they listened to two types of stimuli: Matrix sentences and a natural story. Each stimulus was presented at different levels of speech intelligibility by adding speech weighted noise. Speech intelligibility was assessed in two ways for both stimuli: (1) behaviorally and (2) objectively by reconstructing the speech envelope from the EEG using a linear decoder and correlating it with the acoustic envelope. We also calculated temporal response functions (TRFs) to investigate the temporal characteristics of the brain responses in the EEG channels covering different brain areas.

**Results:** For both stimulus types the correlation between the speech envelope and the reconstructed envelope increased with increasing speech intelligibility. In addition, correlations were higher for the natural story than for the Matrix sentences. Similar to the linear decoder analysis, TRF amplitudes increased with increasing speech intelligibility for both stimuli. Remarkable is that although speech intelligibility remained unchanged in the no noise and +2.5 dB SNR condition, neural speech processing was affected by the addition of this small amount of noise: TRF amplitudes across the entire scalp decreased between 0 to 150 ms, while amplitudes between 150 to 200 ms increased in the presence of noise. TRF latency changes in function of speech intelligibility appeared to be stimulus specific: The latency of the prominent negative peak in the early responses (50-300 ms) increased with increasing speech intelligibility for the Matrix sentences, but remained unchanged for the natural story.

**Conclusions:** These results show (1) the feasibility of natural speech as a stimulus for the objective measure of speech intelligibility, (2) that neural tracking of speech is enhanced using a natural story compared to Matrix sentences and (3) that noise and the stimulus type can change the temporal characteristics of the brain responses. These results might reflect the integration of incoming acoustic features and top-down information, suggesting that the choice of the stimulus has to be considered based on the intended purpose of the measurement.

## 1 INTRODUCTION

In current clinical practice speech intelligibility is measured behaviorally by asking the listeners to recall the words or sentences they heard. By doing so, not only the function of the auditory periphery is measured (Do the speech sounds reach the brain?), but also individual skills like working memory, language knowledge and cognition. When measuring speech intelligibility to evaluate the function of a hearing aid, it is desirable to evaluate the auditory periphery without these extra factors. In addition, the required active participation of the participant can make these measurements challenging or even impossible because of poor attention or motivation, especially in small children.

To overcome these challenges an objective measure of speech intelligibility, where no input from the participant is required, would be of great benefit. Previous studies have shown that the slowly varying speech envelope is essential for speech intelligibility (Shannon et al., 1995), and that it can be reconstructed from brain responses using electroencephalography (EEG) or magnetoencephalography (Luo and Poeppel, 2007; Aiken and Picton, 2008; Ding and Simon, 2011). Correlating the reconstructed envelope from the brain response with the real acoustic envelope, results in a measure of neural envelope tracking, which is related to speech intelligibility in mostly the delta (Ding et al., 2014; Molinaro and Lizarazu, 2017; Vanthronhout et al., 2018) and theta band (Luo and Poeppel, 2007; Ding and Simon, 2013; Lesenfants et al., 2019a).

Vanthornhout et al. (2018) and Lesenfants et al. (2019a) demonstrated the application of this measure of neural envelope tracking in an objective measure of speech intelligibility using isolated Matrix sentences as a stimulus. Matrix sentences are 5-word sentences containing a proper name, verb, numeral, adjective and object with 10 options per word category presented randomly (e.g., ‘Sofie sees ten blue socks’). In their studies the same Matrix sentences were used during a standardized behavioral recall experiment and an EEG measurement, enabling direct comparison of speech intelligibility to envelope tracking. However, for the purpose of clinical applications, the use of isolated sentences may be sub-optimal. Sentences do not reflect everyday communication where syllable, word and sentence rate are less controlled and more semantic top-down processing is involved. Therefore, an objective measure of speech intelligibility based on fully natural speech could (1) overcome subject-related factors such as motivation and therefore possible drops in attention and (2) allow intelligibility measurements of any speech fragment, which is impossible today using behavioral measurements but may relate better to everyday communication.

In this study we investigate whether the objective measure of speech intelligibility by Vanthornhout et al. (2018) using Matrix sentences at a wide range of Signal-to-Noise Ratios (SNRs) can also be conducted with natural running speech, such as a narrated story. We know stories can be used to measure neural envelope tracking (Ding and Simon, 2012, 2013; Kong et al., 2014; Petersen et al., 2017) and Ding and Simon (2013) were able to link neural envelope tracking to speech intelligibility at -3 dB SNR using a story. By replicating the study of Vanthornhout et al. (2018), we want to investigate the relation between neural envelope tracking and speech intelligibility at a wide range of SNRs and compare the use of a natural story to individual sentences to find the most optimal stimulus for clinical use. We hypothesize neural envelope tracking will not be similar for both stimuli as speech intelligibility relies on the active integration of two incoming information streams (Hickok and Poeppel, 2007; Anderson et al., 2018): (1) the bottom-up stream that processes the acoustic features through the auditory pathway until the auditory cortex and (2) the top-down stream originating in different brain regions. Brodbeck et al. (2018) review in their introduction how neural tracking of different speech features has been used to measure top-down and bottom-up processes. We hypothesize that if neural envelope tracking is mainly a feed-forward acoustic process, results for Matrix sentences will be enhanced compared to the story because of the rigid syllable, word and sentence rate reflected in the speech envelope of the Matrix sentences. If, on the other hand, neural envelope tracking captures the interaction between the incoming acoustic speech stream and top-down information, results for the story will be enhanced because of, e.g., increased semantic processing (Di Liberto et al., 2018; Broderick et al., 2018) and attention (Kerlin et al., 2010; Ding and Simon, 2012; Mesgarani and Chang, 2012; Vanthornhout et al., 2019).

## 2 MATERIAL AND METHODS

### 2.1 Participants

Nineteen participants aged between 18 and 28 years (3 men and 16 women) took part in the experiment after providing informed consent. Participants had Flemish as their mother tongue and were all normal-hearing, confirmed with pure tone audiometry (thresholds ≤ 25 dB HL at all octave frequencies from 125 Hz to 8 kHz). The study was approved by the Medical Ethics Committee UZ Leuven / Research (KU Leuven) with reference S57102. All participants were unpaid volunteers.

### 2.2 Auditory stimuli

During the experiment participants listened to three different stimuli: (1) isolated Matrix sentences, (2) a natural story and (3) another story used to train the linear decoder on.

#### 2.2.1 Matrix sentences

Flemish Matrix sentences contain 5 words spoken by a female speaker and have a fixed syntactic structure of ‘proper name-verb-numeral-adjective-object’, for example, ‘Sofie sees ten blue socks’ with a speech rate of 4.1 syllables/second, 2.5 words/second and 0.5 sentences/second. Each category of words has 10 alternatives and each sentence consists of a random combination of these alternatives which induces a rigid and artificial speech rate and reduces semantic context to a bare minimum. These sentences are gathered into lists of 20 sentences. Speech was fixed at a level of 60 dBA and the noise level varied across trials. We used speech weighted noise (SWN) which has the long-term-average spectrum of the stimulus and therefore results in optimal energetic masking. Matrix sentences are a validated speech material to measure speech intelligibility which allows us to directly compare EEG results with speech intelligibility, similar to Vanthornhout et al. (2018) and Lesenfants et al. (2019a). However, Matrix sentences have a rigid speech rate and lack semantic information, resulting in an artificial speech stimulus not representative for everyday communication.

#### 2.2.2 Natural story

The natural story we used is ‘De Wilde Zwanen’, written by Hans Christian Andersen and narrated in Flemish by Katrien Devos (female speaker) with a speech rate of approximately 3.5 syllables/second, 2.5 words/second and 0.2 sentences/second. Speech was fixed at a level of 60 dBA and the noise level of the SWN varied across trials. The main differences between the Matrix sentences (2.2.1) and fully natural speech such as this narrated story are:

1. *Prosody*: Matrix sentences are part of a standardized speech material where every word is spoken at the same intensity, while the story is naturally spoken with intensity variations as a consequence.
2. *Speech rate*: Matrix sentences have a rigid syllable, word and sentence rate, while the story has a naturally varying speech rate because of different word and sentence lengths.
3. *Semantic context*: Matrix sentences are a random combination of words, minimizing the use of semantic context. The story, on the other hand, is coherent speech where the use of top-down processing is triggered, e.g., knowledge about time, space and characters.
4. *Lexical prediction*: The permutations of the words are different in each Matrix sentence, but the words themselves become more familiar to the participants during the experiment, in contrast to the story.

#### 2.2.3 Decoder story

A children’s story, ‘Milan’, written and narrated in Flemish by Stijn Vranken (male speaker), was presented to the participants with a speech rate of 3.7 syllables/second, 2.6 words/second and 0.3 sentences/second. This story is 14 minutes long and was presented at a level of 60 dBA without noise. The purpose of this story was to have an independent continuous stimulus without background noise to train a linear decoder on (Vanthornhout et al., 2018) to reconstruct the speech envelope from the EEG. We intentionally selected a story, because a decoder based on Matrix sentences would be sensitive to sentence onsets, because of the artificially inserted silences, and not to the envelope.

### 2.3 Behavioral experiment

Speech intelligibility was measured behaviorally in order to compare envelope tracking results in terms of speech intelligibility. We need to measure speech intelligibility for both stimuli separately because they differ in content and acoustic parameters (speaker, speech rate, intonation). Adding a similar level of background noise will therefore not result in a similar level of speech intelligibility (Decruy et al., 2018).

Before the EEG experiment we conducted a Matrix test. This test starts with 2 training lists followed by 3 testing lists of 20 sentences each at different SNRs: -9.5; -6.5 and -3.5 dB SNR. Speech was fixed at a level of 60 dB A and the noise level varied in random order. Speech and noise were presented to the right ear. Participants had to recall the sentence they heard. By counting the correctly recalled words, a percentage correct per presented SNR was calculated. Next, a psychometric function was fitted on the data points, similar to what is done in clinical practice.

To measure speech intelligibility for the natural story, we cannot ask the participants to recall every word, instead we used a rating method during the EEG experiment. Participants were asked to rate their speech intelligibility with the following question appearing visually on a screen in front of them: ‘Which percentage of the words did you understand?’ at the presented SNRs (-12.5; -9.5; -6.5; -3.5; -0.5 and 2.5 dB SNR). In addition to the recall procedure for the Matrix sentences before the EEG experiment, we also asked 9 of the 19 participants to rate their speech intelligibility for the Matrix sentences during the EEG, similar to the natural story.

### 2.4 EEG experiment

Ten participants started the EEG experiment by listening to Matrix sentences followed by the natural story. The remaining 9 participants did this in the reversed order. The decoder story was presented in between. The natural story was cut in 7 equal parts of approximately 4 minutes long, which we presented in chronological order. The first part was always presented in silence to optimize comprehension of the storyline. The following 6 parts were presented at 6 different SNRs in random order: -12.5; -9.5; -6.5; -3.5; -0.5 and 2.5 dB SNR. The Matrix sentences were concatenated into 7 lists of 40 sentences with a silent gap between the sentences randomly varying between 0.8 and 1.2 seconds. Each 2-minute trial, containing 40 sentences at a particular SNR, was presented twice to analyze test-retest reliability. The SNRs were the same SNRs as used for the natural story, also in random order and including the condition without noise. To maximize attention and keep the participants motivated, questions were asked about each SNR trial, for example, ‘What happened after sunset?’ (natural story) or ‘Which colors of boats were mentioned?’ (Matrix sentences). The answers were not used for further analysis. After the question, the participants were asked to rate their speech intelligibility with the following question: ‘Which percentage of the words did you understand?’.

### 2.5 Experimental setup

Recordings were made in a soundproof and electromagnetically shielded room. Speech was presented bilaterally at 60 dBA and the setup was calibrated using a 2cm3 coupler of the artificial ear (Brüel & Kjær 4152, Denmark) for each stimulus. The stimuli were presented using APEX 3 (Francart et al., 2008), an RME Multiface II sound card (Germany) and Etymotic ER-3A insert phones (Illinois, USA). A 64-channel BioSemi ActiveTwo (the Netherlands) EEG recording system was used for the EEG recordings at a sample rate of 8192 Hz. Participants sat in a comfortable chair and were asked to move as little as possible during the recordings. There was no fixation point, but they were instructed to keep their eyes open. The participants could see the experimenters screen where questions appeared. They gave their responses by talking through a microphone inside the EEG booth. We inserted a small break between the behavioral and the EEG part and between the Matrix sentences and the natural story if necessary.

### 2.6 Signal processing

In this study we measured neural envelope tracking and linked this to speech intelligibility and stimulus type (natural story versus isolated Matrix sentences). Neural envelope tracking was calculated in two ways: We correlated the acoustic speech envelope (2.6.1) with the speech envelope reconstructed from the EEG response (2.6.2) with the help of a linear decoder. Secondly, we calculated temporal response functions (TRFs) to investigate the temporal characteristics of the brain responses in the EEG channels covering the scalp (2.6.3).

#### 2.6.1 Acoustic envelope

The acoustic speech envelope was extracted from the stimulus according to Biesmans et al. (2017), using a gammatone filterbank followed by a power law. We used a filterbank containing 28 channels spaced by 1 equivalent rectangular bandwidth with center frequencies from 50 Hz until 5000 Hz. The absolute value of each sample in each channel was raised to the power of 0.6. All 28 channel envelopes were averaged which resulted in one single envelope. As a next step, the acoustic speech envelope was band-pass filtered, similar to the EEG signal, in the delta (0.5-4 Hz) or theta (4-8 Hz) frequency band with a Chebyshev filter with 80 dB attenuation at 10% outside the passband. Only these low frequencies were further processed, because they contain the information of interest of the slowly varying speech envelope.

#### 2.6.2 Envelope reconstruction

After applying an anti-aliasing filter, the EEG data was downsampled from 8192 Hz to 256 Hz to reduce processing time and referenced to an average of the electrodes. Next, EEG artefact rejection was done using a multi-channel Wiener filter (MWF) (Somers et al., 2018). the MWF was calculated on the long decoder story without noise and applied on the shorter Matrix and natural story SNR trials. After artefact rejection, the signal was bandpass filtered, similar to the acoustic speech envelope and the sample rate was further decreased from 256 Hz to 128 Hz. A schematic overview is shown in Figure 1.

**Figure 1.**
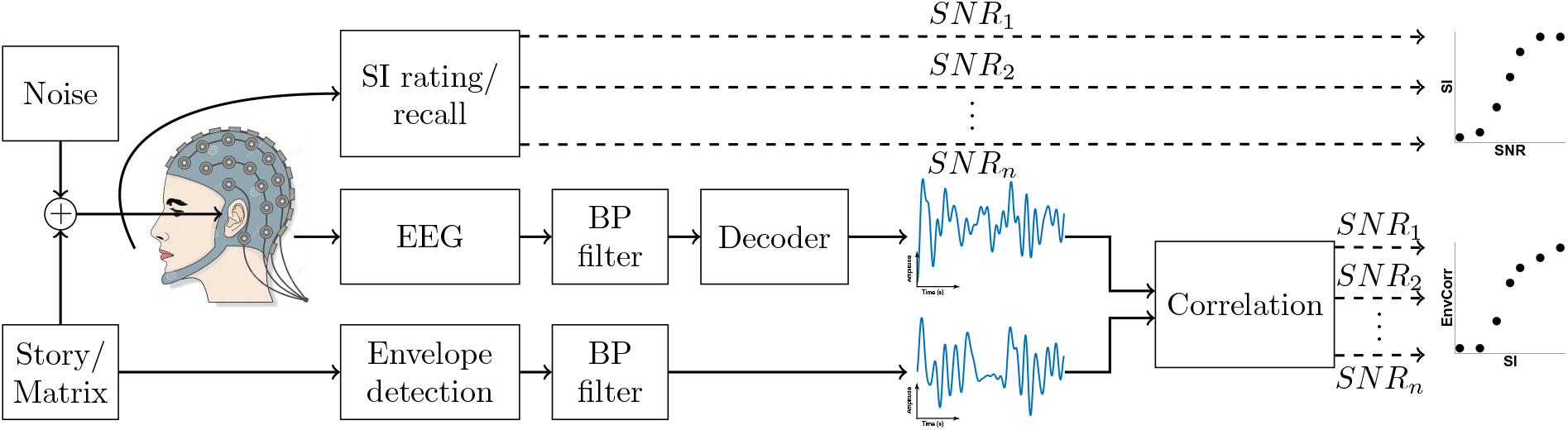
Overview of the experimental setup using the linear decoder analysis. We presented the Matrix sentences and a natural story at different Signal-to-Noise Ratio’s (SNR). Participants listened to the speech while their EEG was measured. To obtain a measure of neural envelope tracking we correlated the reconstructed envelope with the acoustic envelope after band-pass filtering (BP filter). We compared the envelope tracking results with the behavioral speech intelligibility (SI) scores.

To enable reconstruction of the speech envelope from the neural data as a measure of neural envelope tracking, a linear decoder was created in the delta- (0.5-4 Hz) and theta (4-8 Hz) band separately using the mTRF toolbox (Lalor et al., 2006, 2009) similar to the decoder used in Vanthornhout et al. (2018). As speech elicits neural responses with some delay, the decoder not only attributes weights to each EEG channel (spatial filter), but it also takes the shifted neural responses of each channel into account (temporal filter), resulting in a matrix R containing the shifted neural responses of each channel. If *g* is the linear decoder and *R* the shifted neural data, the reconstruction of the speech envelope *s* ^(*t*) was obtained by *s*^ (*t*) = Σ*_n_*Σ*_τ_ g*(*n, τ*)*R*(*t* + *τ, n*) with *t* the time index, *n* ranging over the recording electrodes and *τ* ranging over the integration window, i.e., the number of post-stimulus samples used to reconstruct the envelope. The decoder was calculated by solving *g* = (*RR^T^*)^−1^(*Rs^T^*) with *s* the speech envelope and applying ridge regression to prevent overfitting. We used an integration window of 250 ms post-stimulus resulting in the decoder matrix *g* of 64 (EEG channels) x 33 (time delays within the integration window). The decoder was created using the Milan story (14 minutes) without any noise.

As a last step the envelope was reconstructed by applying the decoder to both test stimuli, the Matrix sentences and the natural story, at various noise levels. Each SNR trial consisted of 2 presentations of 80 seconds of speech after removing the artificially inserted silences between Matrix sentences varying between 0.8 and 1.2 seconds. To measure how similar this reconstructed envelope was to the acoustic envelope as a measure for neural envelope tracking, we calculated the bootstrapped Spearman correlation using Monte Carlo sampling (Pernet et al., 2012) after removing the silences in the stimulus and the corresponding part in the EEG. Removing the silences is necessary as the Matrix sentences contain quasi-regular silent gaps between the sentences which would be a confound. To check whether the obtained correlations were significant, we calculated the significance level of the correlation by correlating random permutations of the real and reconstructed envelope 1000 times and taking percentile 2.5 and 97.5 to obtain a 95% confidence interval. Additionally we calculated the chance levels of both stimuli to investigate whether the decoder has a preference. We hypothesized that because the decoder is a story and not a set of Matrix sentences, the decoder could be better suited to decode the natural story. To obtain the chance levels we reconstructed the envelope of the natural story similar to the standard analysis. Next we correlated the reconstructed envelope of each story trial with the acoustic envelope of all trials of both the natural story (except for the trial used) and the Matrix sentences and compared both.

#### 2.6.3 Temporal response function estimation

The analysis above integrates all neural activity over channels and time lags, i.e. the post-stimulus samples used to create the decoder, and requires a decoder trained on a separate story. To have a closer look at the spatiotemporal profile of the neural responses and remove the assumption that neural processing is similar for the decoder story and the test stimuli in different noise conditions, we calculated TRFs. A TRF is a linear filter that describes how the acoustic speech envelope of the stimulus is transformed into neural responses. It can be used to predict the EEG from the acoustic envelope. This is the inverse approach of the previously mentioned envelope reconstruction where the acoustic envelope is reconstructed from the EEG.

We calculated a TRF for every electrode channel in every participant per stimulus per SNR condition. The first signal processing steps are identical to the envelope reconstruction model starting with downsampling to 256 Hz, artefact rejection with MWF, filtering (0.5-8 Hz) and further downsampling to 128 Hz. Next, TRFs were calculated using the boosting algorithm (David et al., 2007; Brodbeck et al., 2018) with an L2 error norm (using the Eelbrain source code (Brodbeck, 2017)). In summary, boosting is an iterative algorithm starting from a TRF consisting of zeros. With each iteration, the mean-squared error (MSE) is calculated for the prediction after changing all TRF parameters separately by a small amount. The best resulting change after one iteration (smallest MSE), is added to the TRF. This process is repeated until no further relevant improvement is possible (David et al., 2007).

After calculation, the TRFs were convolved with a rotationally symmetric Gaussian kernel of 5 samples long (SD=2) to smooth over time lags. To analyze the TRFs in the time domain, we investigated the latency and amplitude of the negative and positive peaks occurring within 0 and 500 ms after the stimulus (Ding and Simon, 2011; Obleser and Kotz, 2011; Ding and Simon, 2012; Ding et al., 2014).

### 2.7 Statistical Analysis

Statistical analysis was performed using MATLAB (version R2016b) and R (version 3.3.2) software. The significance level was set at *α*=0.05 unless otherwise stated.

For the behavioral tests and envelope reconstruction we compared dependent samples (e.g. test-retest) using a nonparametric Wilcoxon signed-rank test. The correlation between the acoustical envelope and the envelope reconstructed from the EEG was correlated with SI for every filter band and every stimulus using Spearman’s rank correlation. Next, we assessed the relationship between speech intelligibility, envelope reconstruction, filter band and stimulus by constructing a linear mixed effect (LME) model with the following formula:

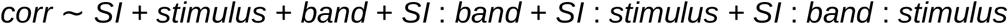

where *corr* is defined as the Spearman correlation between the reconstructed and the acoustic envelope, and fixed and interaction effects of *SI* (speech intelligibility), *stimulus* (Matrix sentences or natural story) and *band* (the delta or theta filter band). An additional random effect of intercept of the participants was included in the model to allow neural tracking to be higher or lower per subject. To control for the different levels of SNRs used for both stimuli to obtain a same level of SI, we constructed the exact same model, but in function of SNR instead of SI.

To control if every chosen fixed and random effect benefited the model the Akaike Information Criterion (AIC) was calculated which estimates a goodness-of-fit measure, while correcting for model complexity. The model with the lowest AIC was selected and its residual plot was analyzed to assess the normality assumption of the LME residuals. Unstandardized regression coefficients (beta) with 95% confidence intervals and p-value are reported in the results section.

To investigate which time samples of the TRF were significantly different from zero, we conducted a cluster-based permutation test. To explore the potential significant differences between the natural story and the Matrix sentences at the different SNRs, we conducted a cluster-based analysis with a post hoc Bonferroni adjustment explained in detail by Maris and Oostenveld (2007). Spearman’s rank correlation was used to investigate the possible change of amplitude and latency of the temporal-occipital peaks over time.

## 3 RESULTS

### 3.1 Behavioral speech intelligibility

During the experiment we measured speech intelligibility behaviorally at different SNRs for every participant. Figure 2 shows that the natural story (rating method) was significantly more difficult than the Matrix sentences (recall method) (p*<*0.001, CI(95%) = [15.99; 23.34], n=19, Wilcoxon signed-rank test). This indicates that the same SNR does not result in the same level of speech intelligibility for the different stimuli. To be able to compare the natural story with the Matrix sentences, we need to account for this.

**Figure 2.**
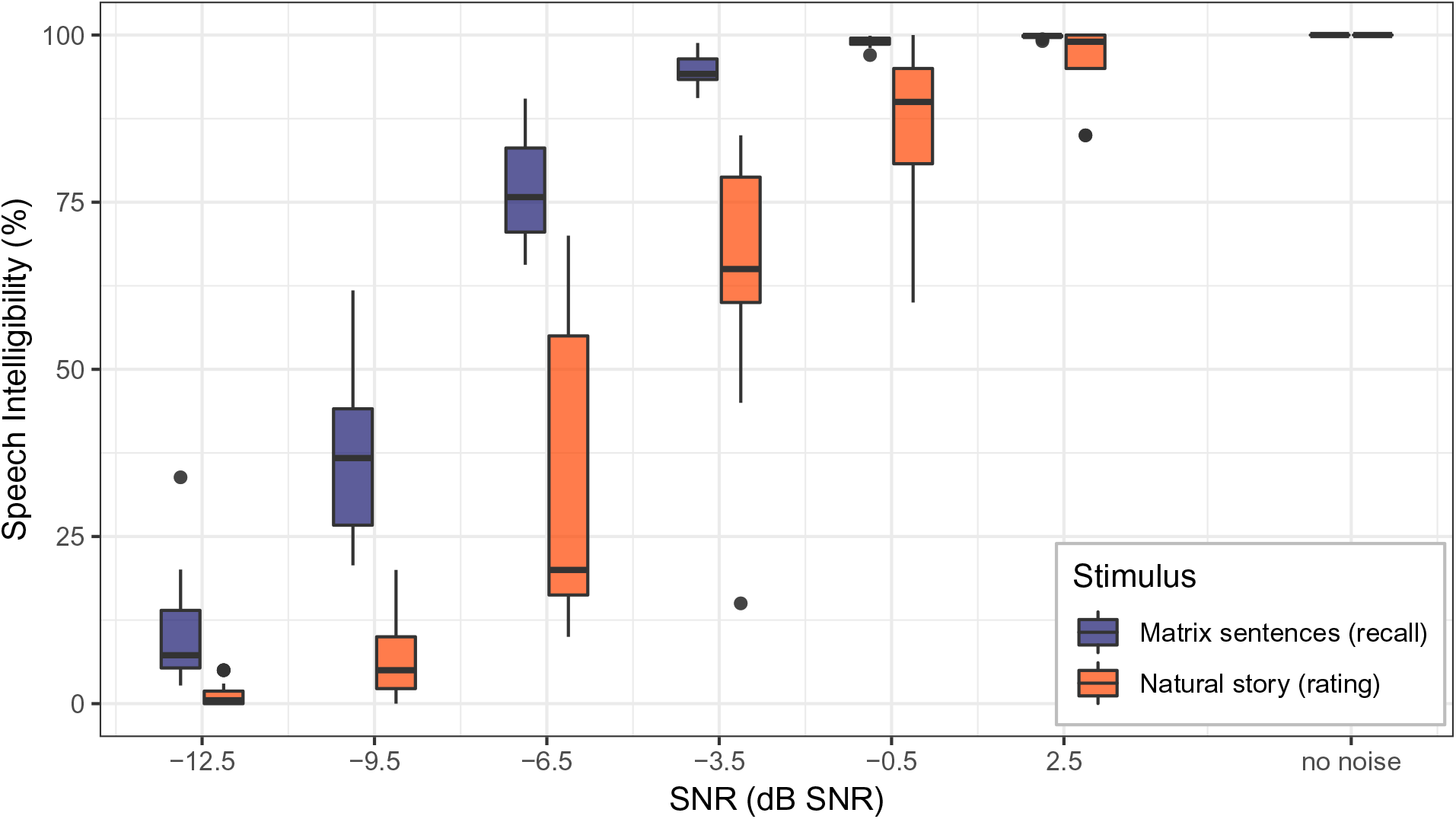
A comparison between the Matrix sentences and the natural story reveals that the story is more difficult to understand when adding background noise.

To check whether the used method to measure speech intelligibility, rate (natural story) versus recall (Matrix sentences), did not influence the results, we asked 9 of the participants to rate their speech intelligibility for the Matrix sentences, similar to the natural story, in addition to the standardized recall method. Comparing their rate and recall scores for the same Matrix sentences at 3 SNRs did not reveal any significant difference (-9.5 dB SNR: p=0.19, CI(95%)=[-11.50; 22.00]; -6.5 dB SNR: p=0.06, CI(95%)=[-29.50; 1.50]; -3.5 dB SNR: p=0.41, CI(95%)=[-9.00; 2.75]; n=9, Wilcoxon signed-rank test).

### 3.2 Envelope reconstruction

To measure neural envelope tracking, we calculated the Spearman correlation between the reconstructed envelope and the acoustic envelope. A test-retest analysis showed no significant difference between test and retest correlations (p=0.746, CI(95%) = [-0.004; 0.006], Wilcoxon signed-rank test), therefore we averaged the correlation of the test and retest conditions resulting in one correlation per participant per SNR per stimulus. Next, no significant difference was found between the chance levels of the stimuli (p=0.534, CI(95%)=[-0.005; 0.003], Wilcoxon signed-rank test). The 95% confidence interval of this non significant difference was similar to the test-retest variability (CI(95%)=[-0.005; 0.006]), indicating that there is no important decoder preference towards one of the stimuli.

We analyzed neural envelope tracking in the delta (0.5-4 Hz) and the theta (4-8 Hz) band for the Matrix sentences and the natural story at various levels of speech intelligibility. Figure 3 shows that when speech intelligibility increases, the correlation between the acoustic and the reconstructed envelope, i.e. neural envelope tracking, increases for every filter band and every stimulus tested (p*<*0.001, table 1, Spearman rank correlation).

**Figure 3.**
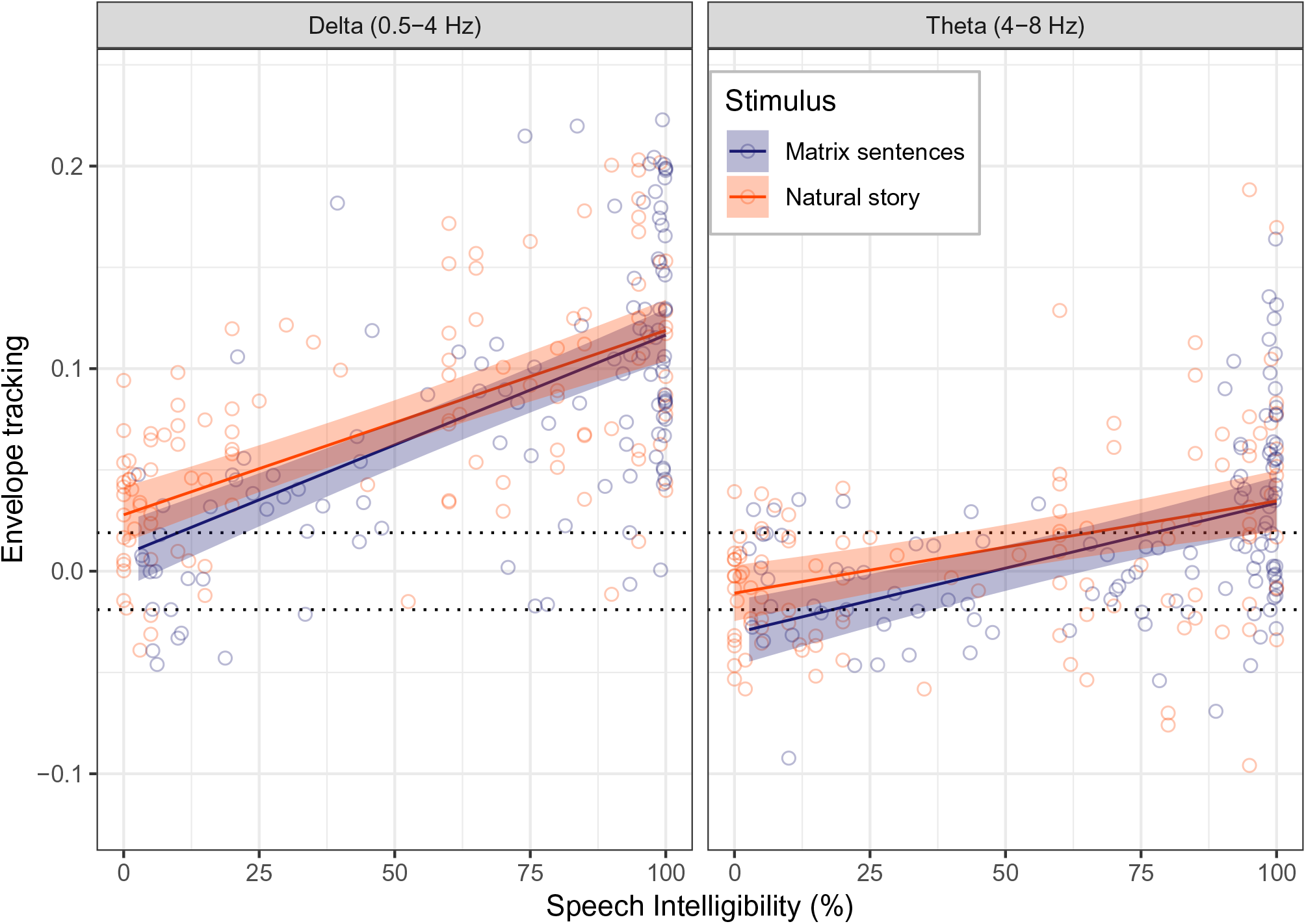
Neural envelope tracking increases with increasing speech intelligibility and by using natural speech as a stimulus. The shading represents two times the standard error of the fit and the dotted line is the significance level of the correlation (+-0.019).

**Table 1.** Spearman rank correlation between neural envelope tracking and speech understanding

| Stimulus | Filter band | Correlation | p-value |
| --- | --- | --- | --- |
| Matrix sentences | Delta (0.5-4 Hz) | 0.62 | p<0.001 |
| Natural story | Delta (0.5-4 Hz) | 0.59 | p<0.001 |
| Matrix sentences | Theta (4-8 Hz) | 0.46 | p<0.001 |
| Natural story | Theta (4-8 Hz) | 0.41 | p<0.001 |

To additionally investigate the influence of stimulus choice, we created an LME model as a function of speech intelligibility. The analysis shows that neural envelope tracking is enhanced for the natural story compared to the Matrix sentences (fixed effect stimulus, p=0.010, LME, table 2). This enhancement does not significantly depend on the level of speech intelligibility or filter band (interaction effect SI:stimulus, p=0.155; interaction effect SI:band:stimulus, p=0.912; LME, table 2). Further, neural envelope tracking in the delta band (0.5-4 Hz) is higher than in the theta band (4-8 Hz) (fixed effect band, p*<*0.001, LME, table 2) with a steeper slope in the delta band (0.5-4 Hz) (interaction effect SI:band, p*<*0.001, LME, table 2).

**Table 2.** Linear Mixed Effect Model of envelope reconstruction in function of SI

| Linear mixed effect model (factor) | beta value | CI(95%) | p-value |
| --- | --- | --- | --- |
| Fixed effect SI | $1.08 \times 10^{-3}$ | $\pm 1.90 \times 10^{-4}$ | $p < 0.001$ |
| Fixed effect stimulus | $1.97 \times 10^{-2}$ | $\pm 1.49 \times 10^{-2}$ | $p = 0.010$ |
| Fixed effect band | $-3.87 \times 10^{-2}$ | $\pm 1.41 \times 10^{-2}$ | $p < 0.001$ |
| Interaction effect SI:stimulus | $-1.74 \times 10^{-4}$ | $\pm 2.39 \times 10^{-4}$ | $p = 0.155$ |
| Interaction effect SI:band | $-4.43 \times 10^{-4}$ | $\pm 2.14 \times 10^{-4}$ | $p < 0.001$ |
| Interaction effect SI:band:stimulus | $-1.28 \times 10^{-5}$ | $\pm 2.25 \times 10^{-4}$ | $p = 0.912$ |

Speech Intelligibility (SI), Confidence Interval (CI)

When conducting the same analysis using SNR as a predictor for speech intelligibility, the same fixed and interaction effects were found to be significant as for the SI analysis (table 3). This shows that even at the same SNR neural envelope tracking for the natural story is enhanced compared to the Matrix sentences, making it impossible to disentangle between the effects of SNR and SI with the current data.

**Table 3.** Linear Mixed Effect Model of envelope reconstruction in function of SNR

| Linear mixed effect model (factor) | beta value | CI(95%) | p-value |
| --- | --- | --- | --- |
| Fixed effect SNR | $7.75 \times 10^{-3}$ | $\pm 1.39 \times 10^{-3}$ | $p < 0.001$ |
| Fixed effect stimulus | $-1.25 \times 10^{-2}$ | $\pm 1.06 \times 10^{-2}$ | $p = 0.022$ |
| Fixed effect band | $-8.10 \times 10^{-2}$ | $\pm 1.06 \times 10^{-2}$ | $p < 0.001$ |
| Interaction effect SNR:stimulus | $-1.01 \times 10^{-3}$ | $\pm 1.83 \times 10^{-3}$ | $p = 0.284$ |
| Interaction effect SNR:band | $-3.20 \times 10^{-3}$ | $\pm 1.83 \times 10^{-3}$ | $p < 0.001$ |
| Interaction effect SNR:band:stimulus | $-1.40 \times 10^{-6}$ | $\pm 2.13 \times 10^{-3}$ | $p = 0.999$ |

Speech-to-Noise Ratio (SNR), Confidence Interval (CI)

### 3.3 Temporal response function

The analysis above integrates all different time lags and channels to obtain an optimal reconstruction of the envelope and requires a decoder trained on a separate story. In the following analysis we focus on how the neural responses follow the envelope in the time and spatial domain and remove the assumption that neural processing is similar for the decoder story and the test stimuli by investigating TRFs. TRFs were calculated on an individual level. This resulted in 868 TRFs per participant (64 channels x 2 stimuli x 7 SNRs). To visualize topographies, we averaged the TRFs per stimulus per SNR over participants. To investigate the time-course of the TRFs, we averaged TRFs for a temporal-occipital channel selection (Figure 4). This selection is based on the TRF results shown in Figure 5. A cluster-based permutation test (Maris and Oostenveld, 2007) shows the TRF samples significantly different from zero, highlighted in bold in Figure 6.

**Figure 4.**
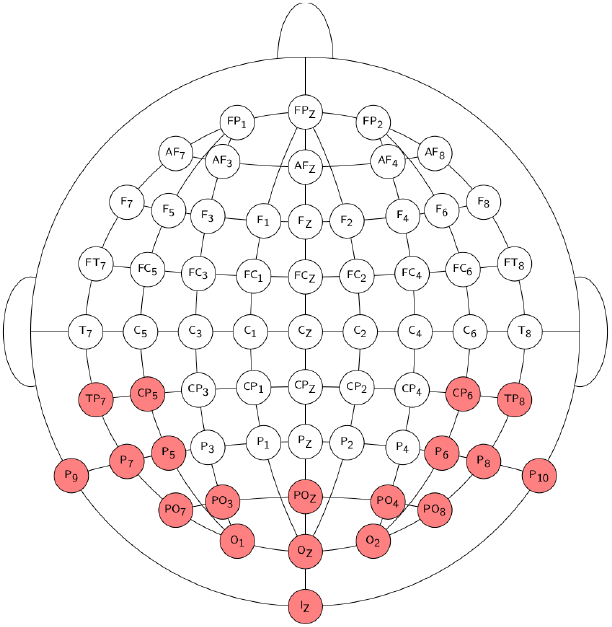
Electrode selection: 64 active electrodes placed according to the 10-20 electrode system. The locations of the electrodes that were selected for the calculation of the occipital-temporal TRF are indicated in red.

**Figure 5.**
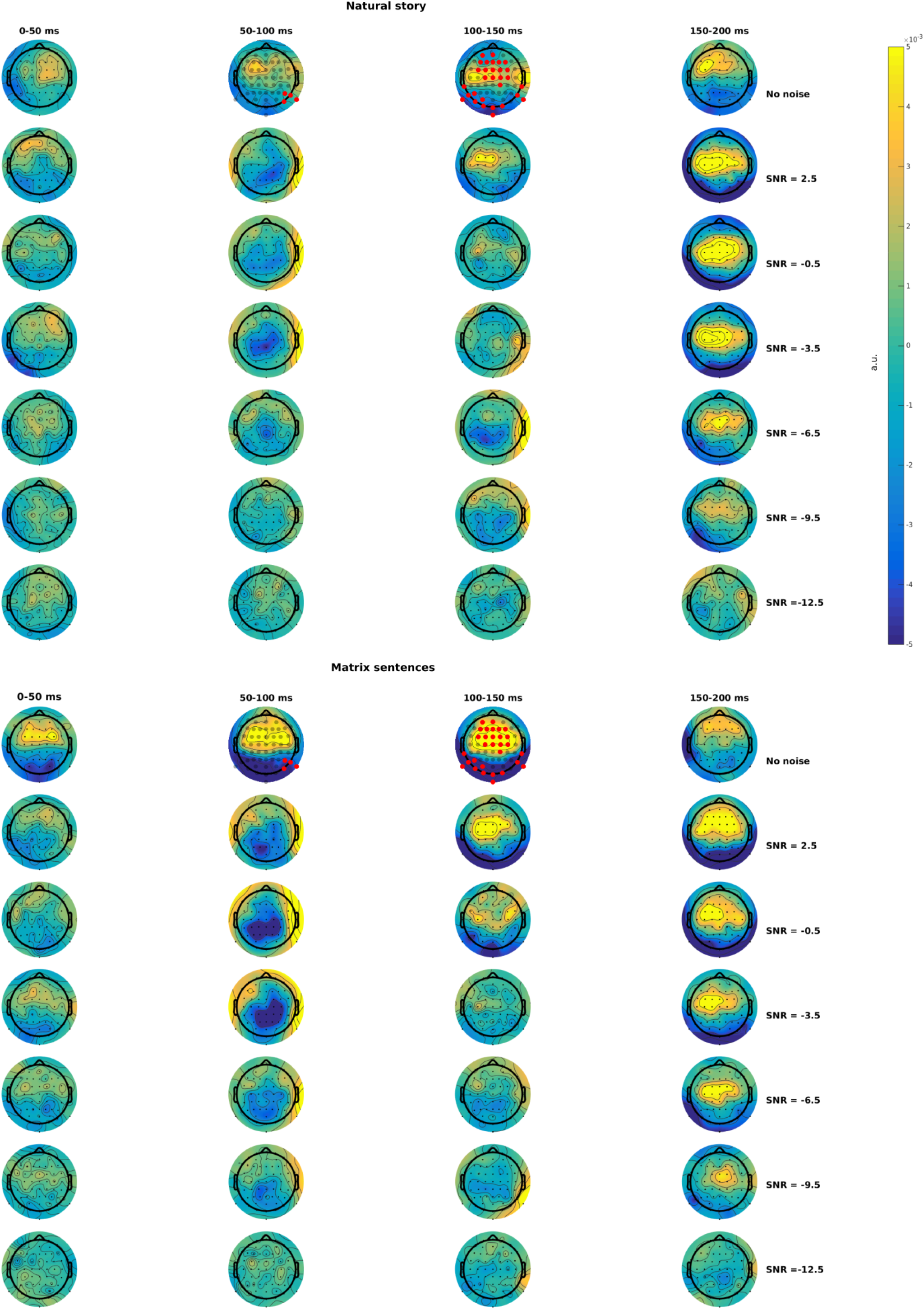
Topographies for the natural story and the Matrix sentences at different SNRs and different time lags varying from 0 until 200 ms. Significant differences between the Matrix sentences and the natural story are highlighted in red.

**Figure 6.**
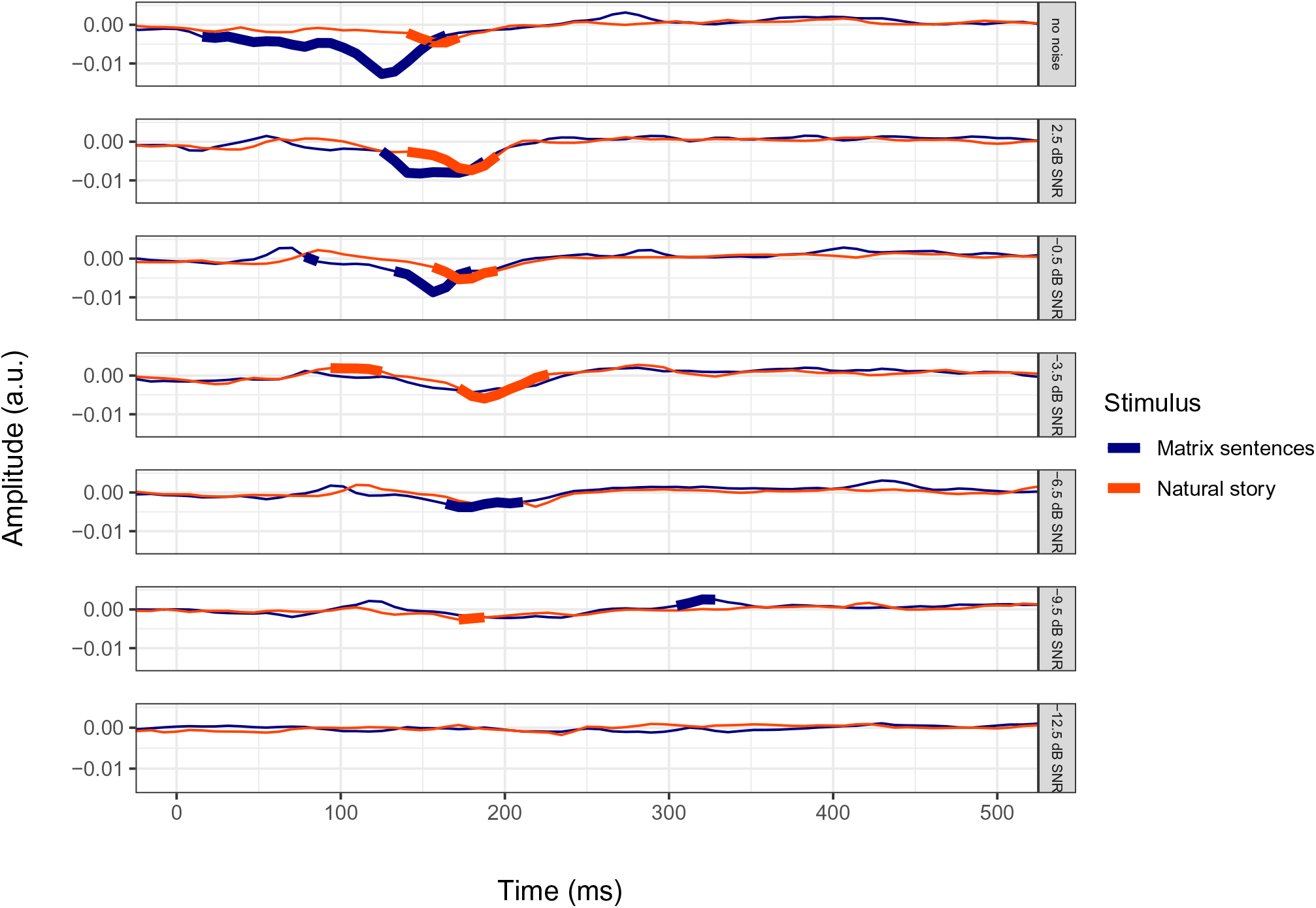
Time-course of the temporal-occipital TRFs over participants for the Matrix sentences and the natural story. TRF samples significantly different from zero are highlighted in bold.

#### 3.3.1 Effect of SNR on TRF

Figure 5 shows the spatiotemporal activation profile of respectively the Matrix sentences and the natural story. In the no-noise condition both stimuli show positive central and negative parieto-occipital amplitudes over time. When a small amount of noise is added and speech intelligibility remains almost unchanged from the no-noise condition (SNR=2.5 dB SNR; Matrix sentences: median SI=99.9%, sd=0.2; Natural story: median SI=99.0%, sd=4.7), the amplitudes across the entire scalp decrease between 0 to 150 ms, while amplitudes between 150 to 200 ms increase in both stimuli. Between 50 and 100 ms amplitudes even swap polarities from positive to negative in the centro-frontal channels (comparing the first 2 rows of the topographies in figure 5). When more noise is added (rows 3-7, figure 5) and speech intelligibility decreases positive central and negative parieto-occipital activation decreases, especially in the 150 to 200 ms time lag. In the 50 to 100 ms time lag, on the other hand, the negative central activation increases with decreasing speech intelligibility and reaches a maximum at SNR=-3.5 dB SNR.

To zoom in on the amplitude changes over time, we visualized an average TRF for the temporal-occipital channels per SNR in Figure 6. When speech intelligibility is very low (SNR<-12.5 dB SNR) both stimuli have very low responses over time. With increasing speech intelligibility, TRF amplitudes also increase gradually. Figure 7 shows the latency and amplitude results of the negative peak that can be found around 100 ms on a participant level over speech intelligibility. It was determined individually by selecting the most negative amplitude of the TRF between 50 and 300 ms. With decreasing speech intelligibility the amplitude of the negative peak per participant decreases for both stimuli (Matrix sentences: Spearman rank correlation=0.49, p*<*0.001; Natural story: Spearman rank correlation=0.26, p=0.005). A centro-frontal channel selection, which is often used in a clinical setting, reveals similar peaks and significances although with different polarity (Appendix A). This is in line with the patterns shown on the topographies in figure 5: Both channel areas are related, the magnitudes vary in the same way over SNR, but the polarities are swapped.

**Figure 7.**
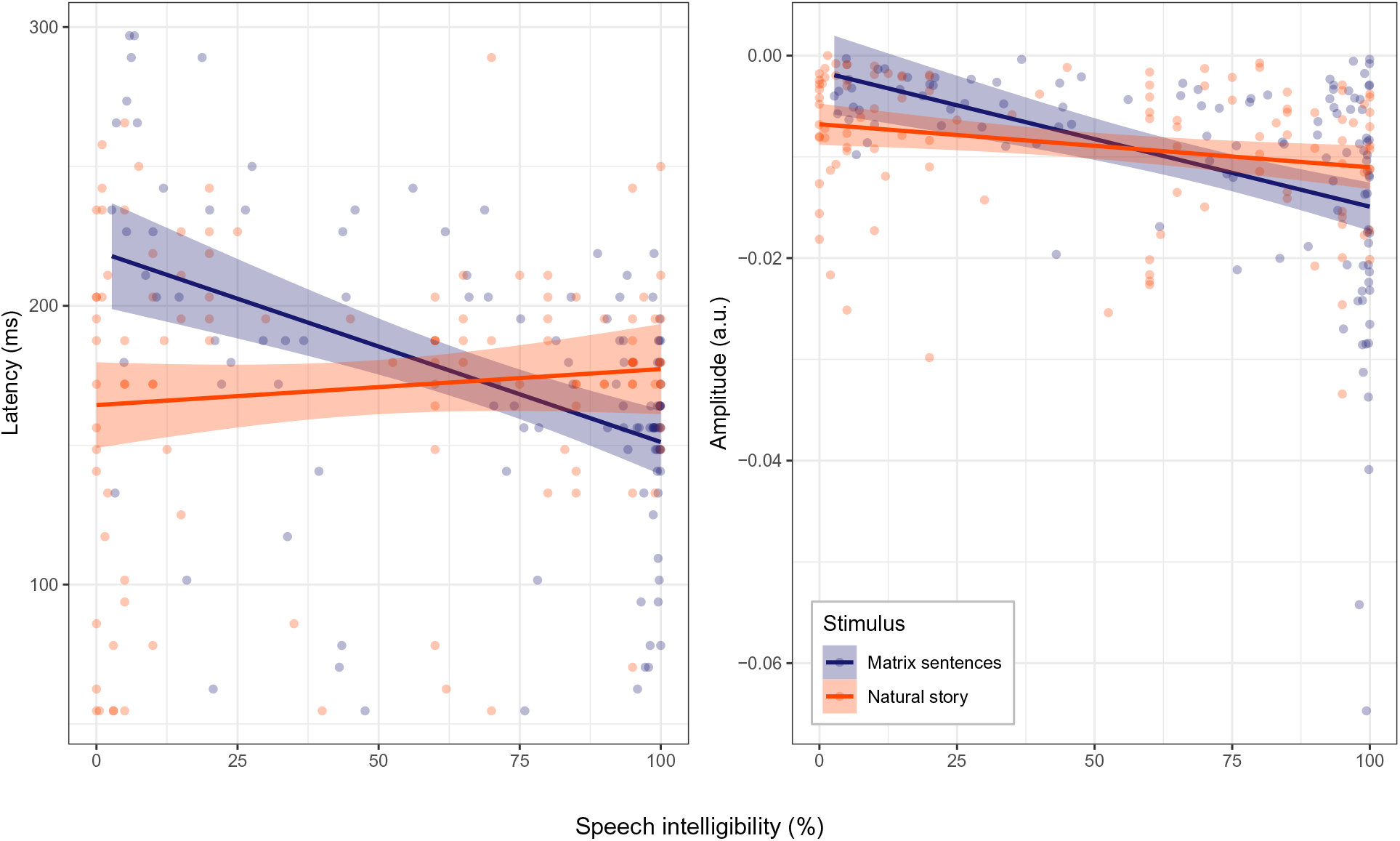
Latency and amplitude of the negative peak of the temporal-occipital TRF between 50 and 300 ms per participant over speech intelligibility.

#### 3.3.2 Effect of stimulus type on TRF

Besides the decreasing amplitude, latency also decreases for the Matrix sentences with decreasing speech intelligibility (Spearman rank correlation=0.46, p*<*0.001). For the natural story, on the other hand, latency is not significantly related to speech intelligibility (Spearman rank correlation=0.02, p=0.835)).

Next to the difference between the Matrix sentences and the natural story concerning latency changes, other stimulus dependent differences can be found. First, a cluster analysis (Maris and Oostenveld, 2007) over all participants revealed significant differences (*α*=0.025) between both stimuli in the no-noise condition with larger amplitudes for the Matrix sentences in the central and parieto-occipital channels, highlighted in red in Figure 5. In contrast to this stimuli driven difference in the no-noise condition, no significant differences between both stimuli could be found in the presence of background noise. Second, in addition to the prominent negative peak between 100 and 200 ms, a positive significant peak arises around 300 ms for the Matrix sentences at -9.5 dB SNR (Figure 6), while this is not the case for the natural story.

## 4 DISCUSSION

In this study we investigated whether the objective measure of speech intelligibility by Vanthornhout et al. (2018) using Matrix sentences can also be conducted with natural speech as this would be beneficial for clinical applications. To that end, we tested 19 normal-hearing participants. They listened to both the Matrix sentences and a natural story at varying levels of speech intelligibility while their EEG was recorded. We found that it is feasible to use natural speech as a stimulus for the objective measure of speech intelligibility and that noise and the stimulus type can change the temporal characteristics of the brain responses over the scalp.

### 4.1 The same SNR does not result in similar speech intelligibility for different stimuli

As a first step we measured speech intelligibility behaviorally for both stimuli at different noise levels. The results show that the same SNR does not result in similar speech intelligibility for the different stimuli. The natural story was found to be more difficult to understand than Matrix sentences. Although we controlled for the sex of the speaker and chose stimuli with similar speech rates and spectrum, the difference could still be due to different acoustic features such as for example prosody. The Matrix sentences namely are part of a standardized speech material where every word is spoken at the same intensity. The natural story, on the other hand, is narrated for children and has more variations. An additional reason to explain this difference is lexical prediction. Even though the permutations of the words are different in each Matrix sentence, the words themselves are all equally likely and familiar to the participants, in contrast to the natural story. Perhaps drawing from a larger pool of words for the Matrix sentences might have led to more similar intelligibility ratings between stimuli. Finally, speech intelligibility for both stimuli was measured in a different way: rating (natural story) versus recall (Matrix sentences). Similar to the very small and insignificant difference of 0.5 dB between rate and recall of Matrix sentences reported by Decruy et al. (2018), we did not find any statistical difference either between both measuring methods applied on the same Matrix sentences.

Besides the difference between the Matrix sentences and the natural story per participant, the variability between participants is also different per stimulus. Variability is high for the natural story compared to the matrix sentences (figure 2). This difference could be explained by the fact that the Matrix sentences are a validated speech material created with the intention to have very low inter-subject variability to only reflect hearing performance independent of other individual skills. The natural story, on the other hand, is not controlled for this and is more dependent on individual skills such as for example linguistic knowledge, cognition and attention span.

### 4.2 Neural envelope tracking as an objective measure of speech intelligibility

We found that the correlation between the reconstructed and the acoustic envelope increased with speech intelligibility for both the Matrix sentences and the natural story. This supports the results of Luo and Poeppel (2007); Ding and Simon (2013); Ding et al. (2014); Molinaro and Lizarazu (2017); Vanthornhout et al. (2018) where an increase in speech intelligibility was also found to accompany an increase in envelope tracking and demonstrates that the objective measure of speech intelligibility using Matrix sentences by Vanthornhout et al. (2018) can be conducted with fully natural speech.

Next, the tracking results in the delta band were significantly higher than in the theta band while the significance levels remain the same, resulting in a steeper slope of envelope tracking as a function of speech intelligibility in the delta band. This difference in correlation magnitude between the frequency bands could be explained by the fact that the modulation spectrum of both stimuli has most energy in the delta band (Luo and Poeppel, 2007; Aiken and Picton, 2008).

When investigating the differences between both stimuli, we found that the use of natural speech enhanced neural envelope tracking compared to Matrix sentences. This suggests that neural envelope tracking might capture the interaction between the incoming acoustic speech stream and top-down information (Hickok and Poeppel, 2007; Gross et al., 2013) such as for example semantic processing (Di Liberto et al., 2018; Broderick et al., 2018). A potential confound is that we used different SNRs for the two stimulus types (to control for intelligibility). This means that the differences in envelope tracking could be related simply to SNR rather than other stimulus properties. To investigate this, we conducted the same analysis, but with SNR as predictor instead of intelligibility, and again found significantly increased envelope tracking for the natural story stimulus. This shows that SNR by itself does not account for the full difference between the two stimulus types. A remarkable finding arising from this SNR analysis, is that neural envelope tracking at a particular SNR is higher for the natural story compared to the Matrix sentences, although SI is lower. This seems in contrast with the hypothesis that neural envelope tracking relates to SI. We hypothesize that this increase is present because the story stimulus elicits more brain activity (e.g., semantic processing and working memory). This underlines the importance of always selecting 1 stimulus to investigate SI to eliminate inter-stimulus differences.

In addition, other confounding factors besides SNR could also be present. First, although the acoustics of the stimuli were matched in terms of sex and speech rate of the speaker and spectrum of the stimulus, acoustic differences like prosody are still present, as discussed in section 4.1. Second, despite the questions asked to motivate the participants, the reduced correlations for the Matrix sentences could be linked to attention. Because listening to concatenated sentences can be boring, attention loss could occur which reduces neural envelope tracking (Ding and Simon, 2012; Kong et al., 2014; Petersen et al., 2017; Vanthornhout et al., 2019). For the natural story, on the other hand, attention could be less of an issue as attending this speech is entertaining possibly resulting in higher correlations.

### 4.3 The effect of noise and stimulus type on neural envelope tracking

In addition to envelope reconstruction to show the feasibility of natural speech as a stimulus for the objective measure (4.2), we conducted a TRF analysis. This analysis enables us to investigate the temporal characteristics of the brain responses over the entire scalp and removes the assumption that neural processing is similar for the decoder story and the test stimuli. The topographies in Figure 5 of both stimuli show a negative activation in the temporal-occipital channels and positive activation in the central channels. This is a typical topography of auditory evoked far-field potentials (Picton, 2011). The large negative peak within the 100 to 200 ms time lag (Figure 6) could be related to the N100, usually occurring at a latency between 70-150 ms (Picton, 2011).

#### 4.3.1 Effect of SNR and speech intelligibility on TRFs

Generally we found, similar to envelope reconstruction, high TRF amplitudes over the entire scalp when speech intelligibility is high (SI=100%) and reduced amplitudes when speech intelligibility decreased for both stimuli, again showing feasibility of natural speech as a stimulus for the objective measure of speech intelligibility. Most remarkable are the TRF amplitudes between 150 to 200 ms, which consistently decrease with decreasing speech intelligibility, perhaps indicating a time window sensitive to speech intelligibility. This amplitude decrease is similar to the behavior of the N1-P2 complex in function of SNR for tone- or syllable-induced event related potentials (ERPs) (Whiting et al., 1998; Billings et al., 2009). Another peculiarity are the noise induced topographic changes. When a small amount of noise is added and speech intelligibility remains almost unchanged from the no-noise condition (SNR=2.5 dB SNR; Matrix sentences: SI=99.9%; Natural story: SI=99.0%), TRF amplitudes across the entire scalp decrease between 0 to 150 ms, while amplitudes between 150 to 200 ms increase. Moreover, TRF amplitudes between 50 and 100 ms even switch polarities in the presence of noise. These increases in amplitudes could potentially be linked to enhanced top-down attention when listening to speech in noise (Fritz et al., 2007). Top-down attention is a selection process that steers neural focusing resources to the desired information stream (the clean speech in this case). This active top-down process causes changes in TRF amplitudes between 50 and 200 ms (Ding and Simon, 2012; Kong et al., 2014; Petersen et al., 2017; Vanthornhout et al., 2019). An additional remark about these noise induced TRF changes concerns the decoder used for envelope reconstruction. Although a decoder trained on clean speech is able to reconstruct the envelope of speech surrounded by noise reliably, it could be more optimal to train the decoder on speech in noise.

#### 4.3.2 Effect of stimulus type on TRFs

Stimulus related differences can be found when comparing topography results between both stimuli. TRF amplitudes are larger for the Matrix sentences in the central and parieto-occipital channels compared to the natural story in the no-noise condition. In the presence of background noise, even at a very high SNR, no significant difference can be found anymore. A possible hypothesis could be the interaction between the incoming acoustic speech stream and top-down information (Hickok and Poeppel, 2007; Gross et al., 2013): In the no-noise condition Matrix sentences are mainly processed in a feed-forward acoustical way. The enhanced TRF amplitudes could be caused by the fixed syntactical 5-word structure of the Matrix sentences, resulting in a more rigid word and sentence rate compared to the natural story. However, when noise is added, more effort has to be paid to listen to the Matrix sentences (Wu et al., 2016; Houben et al., 2013). This changes listening to the Matrix sentences from a bottom-up process to an interactive bottom-up and top-down process similar to the natural story, diminishing the differences between both stimuli.

Another stimulus related difference is the latency pattern over speech intelligibility. The latency of the negative peak decreases with increasing speech intelligibility for the Matrix sentences, while the latency remains unchanged for the natural story. A latency decrease of N100 with increasing speech intelligibility, similar to the Matrix sentences, has been reported in literature analyzing speech- (Petersen et al., 2017; Kong et al., 2014), tone- (Billings et al., 2009) and syllable-induced ERPs (Whiting et al., 1998; Kaplan-Neeman et al., 2006, but is not supported for TRFs for continuous speech by Ding and Simon (2012). This different pattern between the Matrix sentences and the natural story could be explained by two factors. (1) Top-down processing: This is present for the natural story the entire time, for the Matrix sentences, on the other hand, it increases with increasing noise level. Top-down processing requires more time, which could result in delayed TRFs. (2) Attention: Listening to concatenated Matrix sentences might be boring, especially when speech intelligibility decreases, which could result in attention loss and less listening effort known to delay neural processing of speech (Ding and Simon, 2012; Kong et al., 2014; Petersen et al., 2017; Vanthornhout et al., 2019).

A last result to point out is the positive peak around 300 ms for the Matrix sentences at -9.5 dB SNR (SI=49%) (Figure 6). In ERPs a positive peak around this time lag is known to occur when a participant tries to detect a target stimulus (Picton, 1992, 2011). As the Matrix sentences do not contain semantic context, which makes content questions not possible, counting questions were asked at every SNR trial, for example, ‘Which colors of boats were mentioned?’. We hypothesize that the question type, content questions for the natural story versus counting questions for the Matrix sentences, accounts for this difference around 300ms. As a consequence, the type of questions to ask is also an important factor to take into account for future research.

### 4.4 Implications for applied research and potential clinical applications

In this study we showed that the objective measure of speech intelligibility by Vanthornhout et al. (2018) using Matrix sentences can also be conducted with natural speech as a stimulus. Although neural tracking is enhanced using natural speech instead of Matrix sentences, no significant differences in slope of neural tracking in function of speech intelligibility are found. Therefore both stimuli are equally appropriate to calculate the objective measure of speech intelligibility. The stimulus choice will have to be considered based on the intended purpose of the measurement. For clinical applications, for example, we should distinguish between applications in hearing aid fitting and in general diagnostics of the auditory system. For hearing aid fitting, mainly the peripheral processing of speech is of interest, so any type of natural speech is appropriate. For diagnostics, a potential added benefit of a story is that it is closer to everyday communication, and may relate better with subjective experience.

Before clinical application, however, several optimization and validation steps still need to be undertaken: How can we reduce measuring time? How is test-retest reliability within a subject over several test sessions? Also the optimal way of measuring envelope tracking (decoder versus TRF) should be considered. Decoders are probably better suited for clinical applications and TRFs for research. A decoder does not require any channel selection or peak-picking, making it easy to use and interpret the results. TRFs, on the other hand, reveal much more information which is useful for research purposes, but less in the clinical field: Interpreting TRFs is time-consuming and can only be done by highly trained personnel. A concern, however, when using decoders in the clinic, is that subject-specific decoders require long measurement times. A solution would be a generic decoder model which has already been shown to be successful (Di Liberto and Lalor, 2017) and to not decrease performance (Lesenfants et al., 2019b).

### 4.5 Limitations of this study

Frequently recurring confounds throughout the discussion of this study are the attention paid by the listener and listening effort. We tried to control for this by asking the participants to focus and present them content questions after every trial. However, we cannot be sure they always paid attention. In future research it could be useful to also measure attention and listening effort using, e.g., pupillometry (Ohlenforst et al., 2017) or alpha power (Miles et al., 2017; Dimitrijevic et al., 2019). In addition, for the natural story, the order of presented SNRs could also have influenced the results. If, e.g., the higher noise levels were presented first, given that comprehension at the -12.5 SNR level is basically zero, the participant could lose the story line and have worse results in further trials compared to someone who listened to the easier noise levels in the beginning. To minimize this possible bias, we presented all SNRs in random order per participant and the first part of the natural story was always presented in quiet. A third limitation is that the presented Matrix sentences were repeated twice, while the natural story was not. However, we believe this was not a major confound as the Matrix sentences consist of a random combination of 5 word categories which make them very hard to remember. Finally, we rereferenced our data to a common average of the channels to not bias topography patterns relative to the chosen channel. A potential disadvantage of this approach is that the absolute TRF polarity is more difficult to interpret and to compare with previous studies using for example Cz- or mastoid rereferencing. This is, however, not a major concern in this study because we are interested in how TRF polarities and magnitudes of different conditions relate to each other and not to an absolute number.

### 4.6 Conclusion

We found increasing neural envelope tracking with increasing speech intelligibility for both stimuli with an additional enhancement for natural speech compared to Matrix sentences. These results show (1) the feasibility of natural speech as a stimulus for the objective measure of speech intelligibility, (2) that neural envelope tracking is enhanced using a story compared to Matrix sentences and (3) that noise and the stimulus type can change the temporal characteristics of the brain responses.

## Acknowledgements

The authors would like to thank Lien Decruy and Elien Vanluydt for their help in data acquisition. All authors designed the experiment, contributed to the data analysis, discussed the results and implications and commented on the manuscript at all stages. E.V. performed the experiments and wrote the paper. This project has received funding from the European Research Council (ERC) under the European Union’s Horizon 2020 research and innovation programme (grant agreement No 637424 to Tom Francart). Further support came from KU Leuven Special Research Fund under grant OT/14/119. Research of Jonas Vanthornhout (1S10416N) and Eline Verschueren (1S86118N) is funded by a PhD grant of the Research Foundation Flanders (FWO). The authors declare no conflict of interest.

## Appendix A

**Figure 1.**
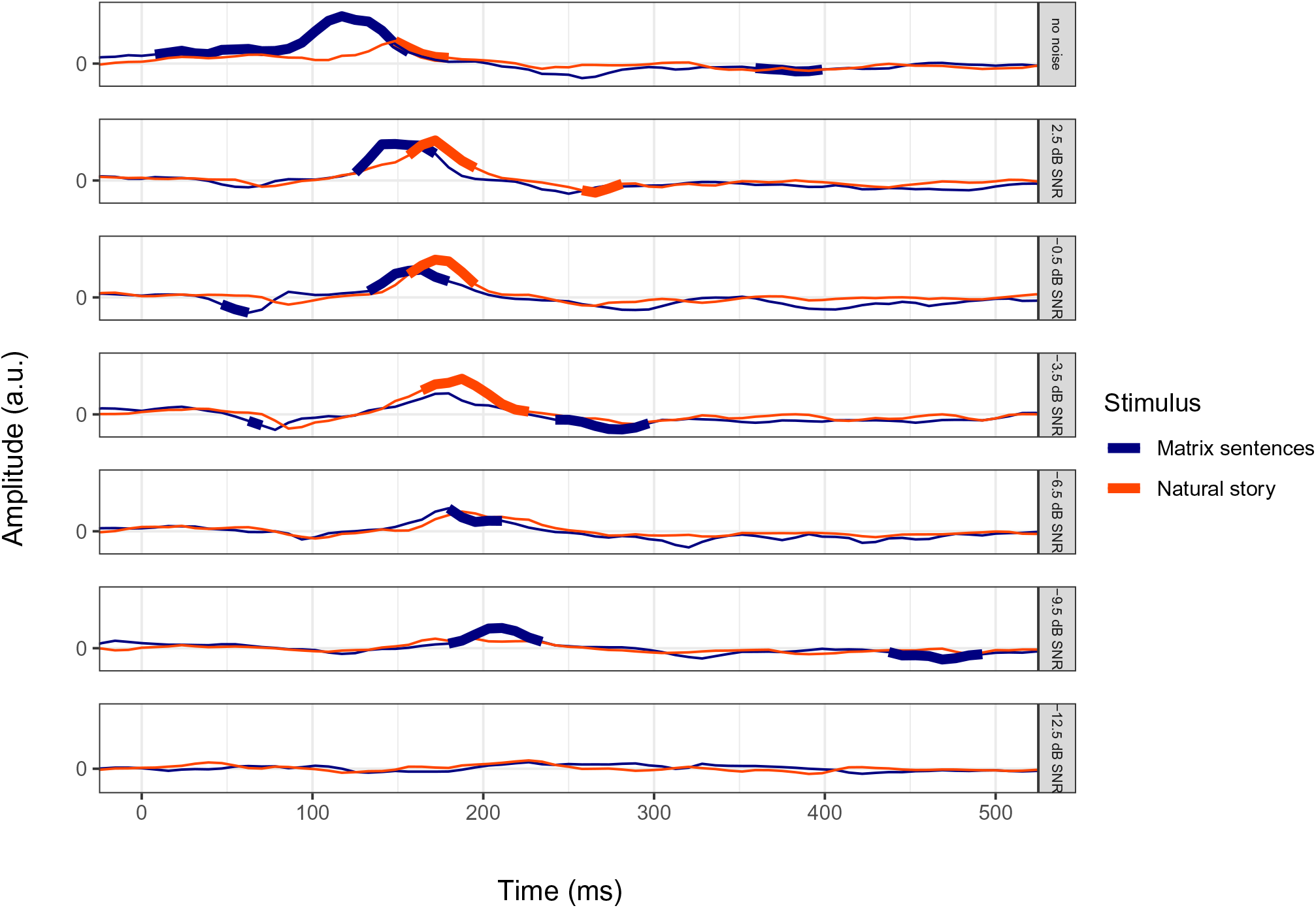
Time-course of the centro-frontal TRFs over participants for the Matrix sentences and the story. TRF samples significantly different from zero are highlighted in bold.

